# Smartphone-Coupled Phase Contrast Microscopy Combined with Deep Transfer Learning for Candida Species Identification: A Proof-of-Concept Study

**DOI:** 10.64898/2026.05.12.724346

**Authors:** Athanasia Sergounioti, Dimitris Rigas, Dimitris Kalles

**Affiliations:** School of Science and Technology, Hellenic Open University, Patras, Greece; Independent Researcher, Amfissa, Greece

**Keywords:** *Candida*, species identification, deep learning, transfer learning, phase contrast microscopy, smartphone microscopy, antifungal therapy, point-of-care diagnostics, resource-limited settings

## Abstract

Species-level *Candida* identification can inform antifungal management, but reliable identification platforms remain inaccessible in many clinical microbiology laboratories, whereas phase contrast microscopy — a common feature of routine laboratory microscopes — is widely available. We asked whether this ubiquitous optical tool, combined with a consumer smartphone and deep transfer learning, could provide a feasible low-cost approach for preliminary *Candida* species discrimination. Fifteen clinical isolates of four species (*C. albicans, C. glabrata, C. tropicalis, C. krusei*) were collected from a single clinical microbiology laboratory and imaged using a consumer-grade smartphone coupled to a standard phase contrast microscope. Suspensions in human serum were imaged immediately after preparation (T0) and after 2-hour incubation at 37°C (T2). Pretrained vision backbone architectures were evaluated as fixed feature extractors under strict Leave-One-Strain-Out cross-validation. The best-performing model — EfficientNet-B0 embeddings with a Linear Support Vector Machine applied to T2 images — achieved an apparent internally cross-validated strain-level balanced accuracy of 0.833 and an overall strain accuracy of 86.7% (13/15 strains correctly classified). *C. albicans, C. glabrata*, and *C. tropicalis* were each identified with 100% recall. Both misclassified strains belonged to *C. krusei* — the species with the smallest panel representation (n=3 strains) — with misclassification attributable to limited strain diversity and suboptimal image quality. These findings demonstrate promising feasibility for preliminary image-based *Candida* species discrimination from smartphone-acquired phase contrast microscopy images, and support further evaluation in larger, externally validated strain collections.

## 1. Introduction

*Candida* species are a leading cause of invasive fungal infections in hospitalised patients, with invasive candidiasis and candidemia associated with substantial crude mortality, commonly reported in the range of 30–60% depending on patient population, species distribution, and clinical context (1–3). Species-level identification is not merely taxonomic: it directly informs antifungal treatment selection, antifungal stewardship, outbreak detection, and surveillance (4,5). *Candida glabrata* and *Candida krusei* exhibit reduced susceptibility or intrinsic resistance to fluconazole, often necessitating alternative agents such as echinocandins or amphotericin B (2,4). *Candida albicans* and *Candida tropicalis* are more commonly azole-susceptible than *C. glabrata* or *C. krusei*, although local susceptibility patterns remain important; species identity can therefore guide early antifungal selection while formal susceptibility testing is pending (2,4,5). Delayed or incorrect species identification may therefore carry direct therapeutic consequences. Beyond antifungal susceptibility, *Candida* species differ in virulence, propensity to form biofilms, and clinical behaviour in immunocompromised hosts — characteristics that further influence management decisions and prognosis (1,2,4). It is this species-level heterogeneity, with its direct bearing on patient outcomes, that sustains ongoing research interest in faster, more accessible, and more accurate identification methods.

Artificial intelligence (AI)-based image analysis applied to clinical microbiology represents an emerging paradigm with the potential to augment labour-intensive identification workflows [6–8]. Deep learning models, particularly convolutional neural networks (CNNs), have demonstrated strong performance in the analysis of microscopy images across infectious-disease applications, including bacterial Gram-stain interpretation, tuberculosis detection, malaria diagnosis, and automated recognition of protozoan or helminth parasites (9–12). Transfer learning — the reuse of feature representations learned on large general-purpose image datasets for specialised downstream tasks — has proven especially effective in biomedical imaging contexts where labelled training data are inherently limited (13,14).

Consumer-grade smartphones now provide high-resolution imaging capabilities and are increasingly used as accessible tools for microscope image digitisation. Smartphone-based microscopy has been explored across point-of-care and low-resource diagnostic contexts, including infectious disease and microbiological applications (15,16). Recent developments in portable optical microscopy, cloud-based image handling, and AI-assisted interpretation further support the feasibility of computationally enhanced microscopy workflows outside highly specialised laboratory environments (17). In parallel, AI-assisted microscopy has recently been applied to Candida-related diagnostic tasks, as summarised in a systematic review of AI-powered microscopic approaches for *Candida albicans* detection, and in primary studies evaluating CNN-based identification of clinically relevant *Candida* species or yeast morphologies in vaginal discharge microscopy (18–20). However, these studies did not evaluate consumer smartphone-coupled phase contrast microscopy combined with serum-induced two-timepoint imaging and strict strain-level transfer-learning validation for *Candida* species discrimination.

To our knowledge, no prior study has evaluated this specific combination of consumer smartphone-coupled phase contrast microscopy, serum-induced two-timepoint imaging, and strict strain-level deep transfer-learning validation for *Candida* species identification. We report a proof-of-concept demonstration that a consumer-grade smartphone coupled to a standard phase contrast microscope, combined with deep transfer learning, can discriminate four clinically relevant *Candida* species with a strain-level balanced accuracy of 0.833 in strict leave-one-strain-out cross-validation. The pipeline encompasses image quality control, dynamic field detection, CLAHE-based preprocessing, systematic CNN backbone screening across ten architectures, and strict strain-level cross-validation. All methodological decisions are reported transparently to enable reproducibility and future extension.

## 2. Materials and Methods

### 2.1 Clinical Isolates

A total of 15 anonymised, non-duplicate *Candida* clinical isolates obtained from different patients during routine diagnostic workup at the Microbiology Laboratory of the General Hospital of Amfissa, Greece, over a one-year period were included in this proof-of-concept study. Isolates were recovered from diverse clinical specimens, including blood cultures, urine, wound swabs, and respiratory samples. No patient-identifiable information was used. Species identification was performed as part of routine laboratory diagnostics using the VITEK 2 Compact automated identification system (bioMérieux, Marcy-l’Étoile, France). The isolate panel comprised four isolates of *Candida albicans* (CA1–CA4), four isolates of *Candida glabrata* (CG1– CG4), four isolates of *Candida tropicalis* (CT1–CT4), and three isolates of *Candida krusei* (CK1–CK3). Clinical isolates were preserved at −86°C in stock culture broth (Bioprepare, Greece) and were subcultured onto Sabouraud dextrose agar before each imaging session, followed by aerobic incubation at 37°C for 24 h.

### 2.2 Image Acquisition

The imaging protocol was adapted from the conventional germ tube test, in which a light yeast suspension is incubated in serum at 35–37°C and examined microscopically after approximately 2 h for germ tube formation (21,22). For each isolate, material from a 24-h Sabouraud dextrose agar subculture was suspended in fresh human serum. Phase contrast microscopy was performed immediately after suspension preparation (T0) and again after 2 h of incubation at 37°C (T2). The two-timepoint design was used to capture both baseline yeast morphology and serum-induced morphological changes.

The biological rationale for the T2 acquisition was that clinically relevant *Candida* species exhibit species-dependent morphological responses after exposure to serum, consistent with the broader capacity of pathogenic *Candida* species to undergo yeast-to-filament transitions under appropriate environmental conditions (21,23). *Candida albicans* typically forms true germ tubes under these conditions, whereas some non-*albicans Candida* species may show pseudohyphal or elongated budding forms. *Candida glabrata*, by contrast, does not produce germ tubes and generally retains a blastoconidial morphology (21–23). These serum-induced differences were intended to provide additional discriminatory morphological information beyond that available at T0.

Images were acquired using a Motorola Moto G72 smartphone with a 108 MP primary sensor and f/1.7 aperture (Motorola, Chicago, IL, USA), coupled to the eyepiece of a Zeiss Axiostar trinocular microscope (Carl Zeiss Microscopy GmbH, Oberkochen, Germany) using a commercial eyepiece-mount smartphone adapter. All images were acquired under phase contrast illumination using a 40× objective. Fields were selected manually, aiming whenever possible to include approximately 50–100 fungal cells per image while avoiding marked overcrowding or debris. Images were saved in JPEG format and organised in a hierarchical folder structure by species, isolate, and timepoint on Google Drive.

Manual smartphone-based acquisition was intentionally retained to reflect a low-cost, non-dedicated imaging setup. This approach introduced some variability in focus, illumination, field alignment, and mechanical stability between imaging sessions, which was subsequently addressed through image quality control and preprocessing. After the image quality-control procedure described below, a mean of 31.6 images per isolate per timepoint was retained for analysis (range, 13–55).

### 2.3 Image Quality Control

All acquired images were subjected to automated quality control (QC) prior to feature extraction. QC was performed using predefined image-level metrics intended to exclude technically inadequate acquisitions while preserving the realistic variability inherent to a smartphone-coupled phase-contrast imaging workflow.

Sharpness was quantified as the variance of the Laplacian of the grayscale image, with images below 18.0 excluded as severely defocused. In-field brightness was computed as the mean pixel intensity within a static circular field mask centered at the image midpoint, with radius equal to 0.34 × min (H, W); images with mean in-field brightness below 45 or above 250 were excluded. The usable signal fraction was defined as the proportion of in-field pixels with intensity values between 8 and 248, and images with usable signal fraction below 0.55 were excluded. An additional artifact indicator was computed as the ratio of mean Sobel-Y to Sobel-X gradient magnitudes; values above 2.2 were recorded as warnings for possible directional acquisition artefacts but were not used as a hard rejection criterion.

A static central circular mask was used for initial QC to provide a simple, reproducible estimate of in-field brightness and usable signal before dynamic field detection was applied during preprocessing (Section 2.4). Images failing any hard-rejection criterion were excluded from all downstream analyses. The QC thresholds were predefined in the analysis pipeline for this smartphone-coupled phase-contrast acquisition setup and were intended to exclude images with major technical limitations, such as severe defocus, marked underexposure or overexposure, and low usable signal, while retaining the realistic image variability expected from a low-cost mobile imaging workflow. The robustness of the main findings to stricter sharpness thresholds was evaluated in sensitivity analyses. These QC thresholds were defined prior to model training and were not optimised against classification performance.

### 2.4 Image Preprocessing

A dynamic circular field detection algorithm was applied to each QC-passed image to delineate the microscope field boundary. The algorithm used Gaussian blurring with a 21 × 21 kernel, thresholding at the 55th intensity percentile with a minimum threshold value of 12, and morphological opening and closing operations with a 15 × 15 kernel to identify the largest valid contour. If no valid contour was detected, a static fallback circular mask centred at the image midpoint was applied, with radius equal to 0.34 × min(H, W), and the fallback event was logged. For each image, mask parameters, including centroid coordinates, area, and detection mode, were stored and used in subsequent preprocessing steps.

Contrast-limited adaptive histogram equalisation (CLAHE) was applied as a fixed preprocessing step to reduce local contrast variation arising from smartphone auto-exposure and inter-session illumination differences. Each RGB image was first converted to grayscale, and pixels outside the detected microscope field were set to zero using the corresponding field mask. CLAHE was then applied to the grayscale image using a clip limit of 2.0 and a tile grid size of 8 × 8 tiles. After CLAHE, the field mask was reapplied so that out-of-field pixels remained zero-valued during CNN feature extraction and downstream analyses. The final processed grayscale image was replicated across three channels to create an RGB-compatible input array for ImageNet-pretrained CNN architectures.

### 2.5 CNN Feature Extraction

Ten ImageNet-pretrained vision backbones were evaluated as fixed feature extractors: EfficientNet-B0, EfficientNet-V2-S, ResNet50, DenseNet121, ConvNeXt-Tiny, Inception-v3, MobileNet-V3-Large, ResNeXt-50-32×4d, Swin Transformer-Tiny, and RegNet-Y-8GF. This transfer-learning strategy was selected because the available dataset was limited in size, comprising 15 clinical strains with approximately 30 quality-control-passed images per strain per timepoint, making end-to-end training of a deep neural network inappropriate because of the high risk of overfitting. ImageNet-pretrained models can provide transferable visual representations for medical image classification tasks, including settings where the target images differ substantially from natural images and where grayscale images are adapted to pretrained RGB-input architectures [24]. Similar deep-learning approaches have recently been explored for Candida and fungal microscopy image analysis, supporting the feasibility of using learned visual representations for yeast/fungal image classification [25,26].

All backbone weights were kept frozen throughout the analysis, and no fine-tuning was performed on the current dataset. For each architecture, the final classification layer was replaced with an identity mapping, so that each processed image was represented as a fixed-length numerical feature vector, hereafter referred to as an embedding. These embeddings represent the image in the high-dimensional feature space learned by the pretrained backbone.

Backbone-specific pretrained transforms, including resizing, centre cropping where required, tensor conversion, and ImageNet normalisation, were applied according to the corresponding torchvision weights configuration. Embeddings were computed separately for T0 and T2 images and stored for subsequent classifier training and strain-level validation.

### 2.6 Backbone and Classifier Selection

Backbone and classifier selection was conducted in two sequential phases to separate the contribution of the pretrained feature extractor from that of the downstream classifier.

In Phase 1, all ten pretrained vision backbones were screened separately for T0 and T2 images using a single fixed downstream classifier: Logistic Regression with balanced class weights. Logistic Regression was selected for this screening phase because it is a simple linear model and therefore introduces limited classifier-specific complexity. Under this design, differences in performance across backbones primarily reflect the discriminative quality of the extracted embeddings rather than the flexibility of the downstream classifier. Because pretrained backbones can vary substantially in transfer performance across downstream tasks and domain-specific datasets, backbone selection was performed empirically within the current dataset rather than assumed a priori (27,28). Performance was assessed under strict leave-one-strain-out cross-validation, as described and justified in Section 2.7. Strain-level balanced accuracy using mean class-score aggregation across held-out images was used as the predefined ranking criterion for backbone selection; image-level metrics were computed as secondary descriptive outputs and were not used for model selection. No prior preference for any backbone architecture was imposed.

In Phase 2, the best-performing backbone–timepoint combination from Phase 1 was carried forward to a systematic comparison of downstream classifier families on the fixed embedding space. Although Logistic Regression provided a controlled screening model in Phase 1, the optimal decision boundary in the high-dimensional embedding space was not assumed to be linear. Different classifier families impose different geometric assumptions on the data; empirical comparison on the extracted embeddings was therefore required. This design is consistent with transfer-learning workflows in which pretrained networks are used to generate learned image representations that are subsequently analysed using conventional machine-learning classifiers (29,30). The following classifiers were evaluated across multiple regularisation or hyperparameter settings: Logistic Regression, PCA-reduced Logistic Regression, Linear Support Vector Machine, Ridge Classifier, Random Forest, and k-Nearest Neighbours. The classifier achieving the highest strain-level balanced accuracy under the same LOSO evaluation framework was selected as the final configuration and was carried forward to the full performance evaluation.

### 2.7 Validation Scheme

Model performance was estimated using strict leave-one-strain-out cross-validation. The biological strain was treated as the unit of cross-validation partitioning and final performance evaluation. In each fold, one complete strain was withheld as the test set, while all images from the remaining strains constituted the training set. Each of the 15 strains contributed exactly one test fold. This grouped validation scheme was used to prevent information leakage at the biological-unit level, since images derived from the same strain cannot be considered statistically independent observations (31,32).

For C. krusei, represented by three strains, each LOSO fold in which a C. krusei strain was withheld retained only two same-species reference strains in the training set. This constraint limits the within-species morphological diversity available for classifier training and is therefore considered when interpreting species-level performance.

Within each fold, StandardScaler normalisation was fitted exclusively on the training-fold embeddings and then applied to the corresponding test-fold embeddings, ensuring that no information from the withheld strain influenced preprocessing.

Strain-level predictions were derived using three aggregation methods. For classifiers without native probability outputs, including LinearSVC, decision scores were converted to probability-like class scores using a softmax transformation prior to aggregation. These scores were used solely for within-strain aggregation and were not interpreted as calibrated probabilities. The primary endpoint was mean class-score aggregation: per-image class-score vectors were averaged element-wise across all images of the held-out strain, and the species with the highest mean score was assigned as the predicted species. Two sensitivity checks were also computed: majority vote, defined as the most frequently predicted class across all images of the held-out strain, and top-k confident mean class-score aggregation, restricted to the half of images with the highest individual confidence scores. Unless otherwise specified, all reported performance figures refer to mean class-score aggregation.

Bootstrapped 95% confidence intervals for strain-level balanced accuracy were computed using 1,000 bootstrap iterations with strain-level resampling with replacement from the LOSO prediction table. This prediction-level bootstrap approach approximates sampling variability in the observed strain-level prediction outcomes without requiring model retraining. Accordingly, these intervals are interpreted as approximate descriptive intervals rather than fully unbiased estimates of generalisation uncertainty.

## 3. Results

### 3.1 Image Quality Control

Of 1,047 acquired images across all strains and timepoints, 948 passed quality control and were included in downstream analysis, corresponding to an overall pass rate of 90.5%. All T0 images passed QC criteria (476/476, 100%). In contrast, 472 of 571 T2 images passed QC (82.7%), with low sharpness representing the principal reason for image rejection.

QC rejection at T2 was heterogeneous across strains and species. Among the T2 images, C. krusei retained 82/94 images (87.2%), whereas C. tropicalis retained 83/129 images (64.3%), representing the highest T2 rejection burden among the analysed species. The elevated T2 rejection rate in C. tropicalis was consistent with acquisition variability across imaging sessions rather than a systematic failure of species-level image acquisition; importantly, all four C. tropicalis strains were correctly classified in the downstream strain-level analysis, indicating that the retained QC-passed images preserved sufficient discriminative information.

Within C. krusei, image-quality loss was concentrated in strain CK2. For this strain-timepoint combination, 12 of 25 raw T2 images were rejected, leaving 13 QC-passed images available for analysis. The mean sharpness of QC-passed CK2 T2 images was substantially lower than that of CK1 T2 images, whereas sharpness values across the remaining species were within the expected range for smartphone-coupled phase-contrast acquisition. This quality differential was subsequently considered when interpreting the lower downstream classification performance observed for C. krusei. Dynamic field detection was successfully applied to all 948 QC-passed images, and no fallback static mask was required.

### 3.2 Backbone Screening

Screening of ten pretrained vision backbones across two timepoints, corresponding to 20 backbone– timepoint combinations, identified EfficientNet-B0 applied to T2 images as the top-performing feature extractor. This combination achieved a strain-level balanced accuracy of 0.708 and an overall strain accuracy of 0.733 under LOSO evaluation. T2 embeddings generally outperformed T0 embeddings across the screened backbones, yielding higher strain-level balanced accuracy in seven of ten architectures. ResNet50 and Swin-T showed slightly higher performance at T0, while MobileNet-V3-Large showed equivalent strain-level balanced accuracy at both timepoints. The five highest-ranking backbone–timepoint combinations from Phase 1 screening are shown in Table 1.

**Table 1.**
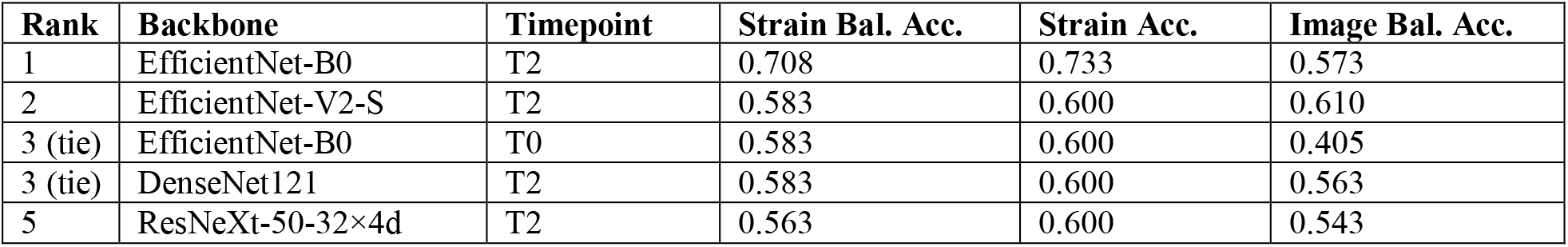
Top five backbone–timepoint combinations from Phase 1 screening, ranked by strain-level balanced accuracy under mean class-score aggregation using leave-one-strain-out cross-validation. Bal. Acc. = balanced accuracy.

### 3.3 Classifier Tuning and Final Performance

Classifiers from six families were evaluated across predefined hyperparameter grids on the fixed EfficientNet-B0/T2 embedding space under the same LOSO framework; results are summarised in Table 2. LinearSVC with C=0.1 achieved the highest strain-level balanced accuracy (0.833), outperforming Logistic Regression (best: 0.771 at C=5.0), Ridge Classifier (best: 0.708 at α=10.0), k-Nearest Neighbours (best: 0.708 at k=3), PCA-reduced Logistic Regression (best: 0.563) and Random Forest (0.563). The performance advantage of LinearSVC was consistent across regularisation settings, providing a clear empirical basis for its selection as the final classifier.

**Table 2.** Phase 2 classifier comparison on the fixed EfficientNet-B0/T2 embedding space under leave-one-strain-out cross-validation. Best configuration per family shown. Bal. Acc. = balanced accuracy (mean class-score aggregation).

| Classifier | Strain Bal. Acc. | Strain Acc. |
| --- | --- | --- |
| **LinearSVC ($C=0.1$)** | **0.833** | **0.867** |
| Logistic Regression ($C=5.0$) | 0.771 | 0.800 |
| Ridge Classifier ($\alpha=10.0$) | 0.708 | 0.733 |
| k-Nearest Neighbours ($k=3$) | 0.708 | 0.733 |
| PCA-LR (100 components) | 0.563 | 0.600 |
| Random Forest | 0.563 | 0.600 |

Under LOSO evaluation, this configuration achieved a strain-level balanced accuracy of 0.833 and an overall strain accuracy of 0.867, with an approximate bootstrapped 95% confidence interval of 0.750–1.000 for strain-level balanced accuracy. Mean class-score aggregation and top-k confident mean class-score aggregation produced identical strain-level results. In contrast, majority vote yielded a substantially lower balanced accuracy of 0.646, indicating that score-based aggregation was more effective than vote-based aggregation in this small strain-level panel. Per-species performance of the final configuration is presented in Table 3.

**Table 3.**
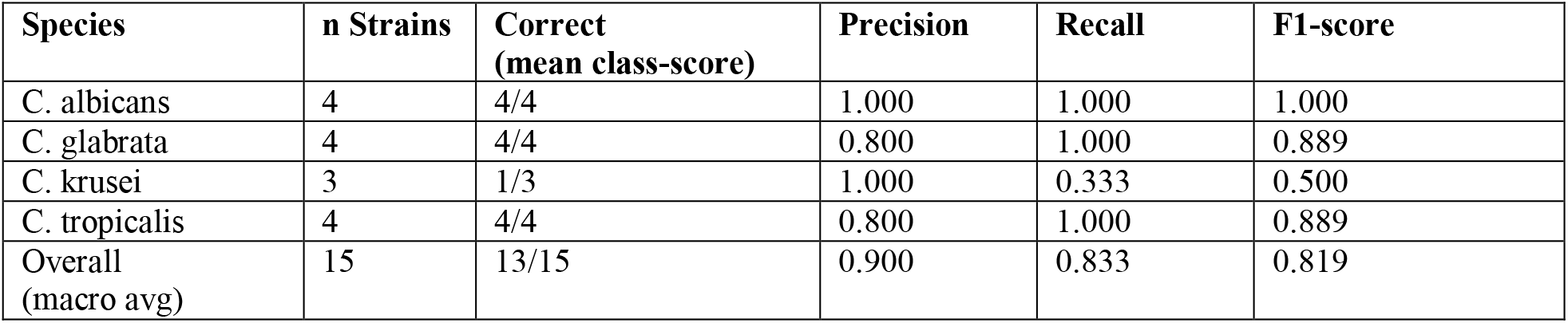
Per-species strain-level classification performance of the final EfficientNet-B0/T2 + LinearSVC model under leave-one-strain-out cross-validation. Metrics were computed at the strain level using mean class-score aggregation.

| Species | n Strains | Correct<br>(mean class-score) | Precision | Recall | F1-score |
| --- | --- | --- | --- | --- | --- |
| *C. albicans* | 4 | 4/4 | 1.000 | 1.000 | 1.000 |
| *C. glabrata* | 4 | 4/4 | 0.800 | 1.000 | 0.889 |
| *C. krusei* | 3 | 1/3 | 1.000 | 0.333 | 0.500 |
| *C. tropicalis* | 4 | 4/4 | 0.800 | 1.000 | 0.889 |
| Overall<br>(macro avg) | 15 | 13/15 | 0.900 | 0.833 | 0.819 |

*C. albicans, C. glabrata*, and *C. tropicalis* each achieved perfect strain-level recall (1.000). The precision of 0.800 for *C. glabrata* and *C. tropicalis* reflected one false-positive assignment to each species, both originating from misclassified *C. krusei* strains: CK2 was assigned to *C. tropicalis*, and CK3 was assigned to *C. glabrata*. These errors indicate that the main residual classification difficulty was concentrated in *C. krusei*. From a susceptibility-pattern perspective, the CK2 assignment to C. tropicalis would be the more concerning misclassification; confusion between C. krusei and C. glabrata may be less divergent with respect to expected fluconazole activity. However, no treatment inference should be drawn from these assignments in the pipeline’s current form.

The softmax-transformed class score assigned to CK3 as *C. glabrata* was high (0.741), whereas the only correctly classified *C. krusei* strain, CK1, had a lower winning class score (0.323), consistent with weaker class separation for *C. krusei*. Image-level balanced accuracy was substantially lower than strain-level balanced accuracy (0.577 vs. 0.833), underscoring the importance of strain-level aggregation when individual image predictions are noisy.

### 3.4 *C. krusei* Misclassification Analysis

To further investigate the *C. krusei* misclassifications, sharpness distributions of QC-passed T2 images were compared across strains. CK1, the only correctly classified *C. krusei* strain, showed a T2 sharpness distribution comparable to well-focused strains from the other species. In contrast, CK2 had a median T2 sharpness close to the QC rejection threshold, with 12 of 25 raw T2 images excluded during QC. CK3 showed intermediate sharpness values.

Although CK2 and CK3 were misclassified, their Laplacian sharpness distributions were not uniformly lower than those of all correctly classified non-*C. krusei* strains. For example, the median T2 sharpness for *C. tropicalis* was 22.0, and several correctly classified *C. albicans* and *C. tropicalis* strains had median sharpness values below 30. This suggests that the Laplacian-based QC metric captured gross image degradation but did not fully reflect all aspects of focus quality relevant to downstream classification. Visual inspection of CK2 and CK3 T2 images supported this interpretation, indicating that some borderline-focus images were retained despite suboptimal visual quality.

A second contributing factor was the small *C. krusei* panel. Because only three *C. krusei* strains were available, each LOSO fold testing a *C. krusei* strain was trained on only two same-species reference strains. The combination of limited within-species training diversity and borderline image quality provides the most parsimonious explanation for the concentration of errors in *C. krusei*.

## 4. Discussion

This proof-of-concept study provides preliminary evidence that consumer-grade smartphone-coupled phase-contrast microscopy, when combined with deep transfer learning and rigorous strain-level cross-validation, can support preliminary *Candida* species discrimination in a clinical microbiology setting. The final pipeline achieved an apparent internally cross-validated strain-level balanced accuracy of 0.833 and an overall strain accuracy of 0.867, with perfect recall for *C. albicans, C. glabrata*, and *C. tropicalis*. These species include major clinically relevant causes of invasive *Candida* infection, for which species-level identification can inform early antifungal management and stewardship (1–5). Importantly, the workflow was implemented using a standard phase-contrast microscope coupled to a consumer-grade smartphone, supporting the feasibility of a low-cost image-based approach in laboratories where advanced identification platforms may be less accessible (15–17).

EfficientNet-B0 was the best-performing feature extractor in the present screening, outperforming larger or more complex architectures such as ResNet50, DenseNet121, and Inception-v3 on the predefined strain-level ranking criterion. This finding supports the need for empirical backbone selection in small, domain-specific image datasets, rather than assuming that larger or more complex architectures will necessarily perform better. Transfer-learning studies in medical imaging show that pretrained models can provide useful image representations in limited-data settings (13,14), while recent backbone-comparison studies demonstrate that transfer performance varies substantially across downstream tasks and domain-specific datasets (27,28). The systematic screening conducted in this study, encompassing ten pretrained vision backbones across both timepoints without any prior architecture preference, therefore provided an empirically grounded and transparent basis for model selection.

The overall performance advantage of T2 over T0 images in most screened backbones is biologically interpretable. Incubation in human serum at 35–37°C is the basis of the conventional germ tube test and promotes time-dependent morphological changes in Candida, most notably germ-tube and filamentous growth in C. albicans (21,22). Recent work further supports a mechanistic role for serum components, particularly albumin, in modulating C. albicans germ-tube formation and filamentation (33). More broadly, pathogenic Candida species can exhibit morphology-associated phenotypic transitions, although the extent and nature of these responses differ substantially between species (23). In parallel, recent image-analysis work on C. albicans filamentation highlights the biological and computational relevance of microscopy-based quantification of serum- or host-cue-associated morphological changes (34). These serum-induced and time-dependent morphological differences may increase inter-species separability in the pretrained embedding space, explaining the observed advantage of T2 embeddings despite greater acquisition variability at this timepoint.

The final model also highlighted the importance of strain-level aggregation. Mean class-score aggregation and top-k confident mean class-score aggregation produced identical strain-level performance, whereas majority vote performed substantially worse. This indicates that, in this small image panel, retaining relative class-score information across all images of a held-out strain was more informative than relying only on the most frequent image-level label. Conceptually, this resembles multiple-instance learning settings, in which a higher-level label is inferred from a collection of lower-level image instances rather than from a single image alone (35,36). In the present study, this principle was implemented as a simple deterministic aggregation step rather than as a trainable multiple-instance learning model.

Both misclassified strains belonged to *C. krusei*. Failure-mode analysis identified two complementary contributors. First, visual inspection indicated that CK2 and CK3 T2 images had borderline or suboptimal focus quality that was not fully captured by the Laplacian-based QC filter, highlighting an operational limitation of manual smartphone-based acquisition. CK1, whose T2 images were visually better focused, was correctly classified, suggesting that the pipeline can retain discriminatory information for *C. krusei* when image quality is adequate. Second, the *C. krusei* panel was small, with only three strains available. Consequently, each LOSO fold testing a *C. krusei* strain was trained on only two same-species reference strains, limiting the within-species morphological diversity available to the classifier. The combination of borderline image quality in two of the three *C. krusei* strain–timepoint sets and minimal same-species training diversity provides the most parsimonious explanation for the observed concentration of errors in *C. krusei*.

This study has several limitations inherent to its proof-of-concept design. The strain panel was small, comprising 15 strains across four species, and was unbalanced, with *C. krusei* represented by only three strains. Because backbone and classifier selection were performed on the same small strain panel used for internal LOSO performance estimation, the reported performance should be interpreted as an apparent internally cross-validated estimate and may be optimistic. Larger external validation cohorts will be required to obtain less biased estimates of diagnostic performance. The bootstrapped confidence intervals reported here were derived from prediction-level resampling without model retraining and should therefore be interpreted as approximate descriptive intervals rather than fully unbiased estimates of generalisation uncertainty.

Species identification was based on routine laboratory assignment using VITEK 2 Compact rather than MALDI-TOF MS or molecular confirmation. Reference-label uncertainty therefore cannot be fully excluded, and the present study should be interpreted as a feasibility assessment relative to routine laboratory species assignment rather than as definitive diagnostic validation against a reference-standard identification method. However, the documented accuracy limitations of VITEK 2 are most pronounced for uncommon and cryptic *Candida* species absent from or underrepresented in its reference database; for the four species comprising the present panel, concordance between VITEK 2 and MALDI-TOF MS has been reported to be high in comparative studies (37,38). Species assignment for *C. krusei* isolates was additionally corroborated by characteristic colonial morphology — flat, dry, and distinctly matt surface texture — which is phenotypically inconsistent with the other three species in the panel, providing independent macroscopic support for the reference labels. MALDI-TOF MS and sequencing-based approaches provide more robust species-level identification for yeasts, particularly where species-complex or cryptic-organism resolution is required (39,40). Future validation cohorts should therefore use MALDI-TOF MS and/or molecular confirmation as reference standards. The imaging setup, based on manual smartphone coupling, produced variable image quality across acquisition sessions; a fixed digital microscope camera connected via the trinocular port, or a mechanically standardised smartphone adapter with robust calibration, would be expected to improve consistency, in line with recognised requirements for smartphone-based biomedical imaging systems (41). Finally, all isolates were collected from a single centre, which may limit the generalisability of the learned representations to strains from geographically or epidemiologically distinct populations.

Future work will focus on systematic expansion and external validation of the pipeline. The immediate priority is to increase the number of strains per species, with species identification confirmed by MALDI-TOF MS and/or molecular methods as reference standards. Lower-quality strain–timepoint combinations, particularly CK2 and CK3 at T2, should also be reacquired under improved imaging conditions to assess whether the observed *C. krusei* errors are reproducible or primarily acquisition-related. In subsequent work, the pipeline should be evaluated on additional clinically relevant *Candida* species, including *C. parapsilosis, C. dubliniensis*, and *C. auris*. The latter is of particular interest because of its emerging multidrug-resistant profile and the documented challenges of reliable identification by some conventional laboratory systems (42). Cross-instrument validation across different microscopes, cameras, and smartphone models, together with multicentre validation using geographically and epidemiologically diverse isolates, will be essential before the generalisability and potential clinical utility of the approach can be assessed.

From a practical standpoint, the proposed workflow is simple and low-cost. When fungal growth is observed on primary culture, a yeast suspension can be prepared in human serum and imaged immediately at T0 and again after approximately two hours of incubation at 37°C. Phase-contrast microscopy images can then be acquired using a smartphone coupled to a standard microscope, and the computational pipeline can generate a species-level prediction within minutes of T2 image acquisition. In principle, this could provide a same-working-day preliminary indication after first growth detection, without additional subculture or prolonged incubation beyond the short serum-incubation step. At the current stage of development, however, the pipeline output should be regarded only as a preliminary, non-standalone image-based indication requiring confirmation by established identification methods.

In conclusion, this proof-of-concept study demonstrates that smartphone-coupled phase-contrast microscopy combined with deep transfer learning can provide a feasible preliminary image-based approach for *Candida* species discrimination. The pipeline is methodologically transparent, intended to be released as open-source upon acceptance, and designed for extension to larger, more diverse, and externally validated strain panels. These findings support further investigation of the approach as a low-cost, non-standalone preliminary indication requiring confirmation by established laboratory identification methods before any clinical use can be considered.

## Declarations

### Ethics approval and consent to participate

Not applicable. Isolates were obtained as part of routine diagnostic workup; no patient-identifiable data were collected or used.

### Availability of data and materials

The analysis pipeline and image dataset are available from the corresponding author upon reasonable request.

### Authors’ contributions

AS conceived the study, designed the experimental protocol, acquired the microscopy images, contributed to the design and structuring of the computational pipeline, interpreted the findings, and wrote the original manuscript draft. DR Dimitris Rigas developed and implemented the computational pipeline, including image preprocessing, deep feature extraction from pretrained vision backbones, model training, leave-one-strain-out validation, performance analysis, and contributed to the interpretation of the results. DK supervised the study, provided methodological and scientific guidance, and critically revised the manuscript. All authors read and approved the final manuscript.

